# Spatial separation of *Streptomyces* aerial mycelium during fermentation enhances secondary metabolite production

**DOI:** 10.1101/2020.05.27.118091

**Authors:** Balaji Muralikrishnan, Retnakumar R J, Ranjit Ramachandran, Vinodh J S, Nijisha M, Arun K B, Laiza Paul, Syed Dastager, Vipin Mohan Dan, Ajay Kumar Ramakrishnan

## Abstract

Conditioning of morphology is an effective technique to enhance secondary metabolite production by *Streptomyces*. Here we report a novel conditioning method employing glass marbles in batch cultures to enhance secondary metabolite production by *Streptomyces sp*. The marbles seem to spatially separate aerial and submerged mycelia in the flask which was necessary for the qualitative and quantitative enhancement of metabolite production of secondary metabolites. The method also offers shorter incubation period compared to conventional methods for effective production. Further, using a combination of this method and response surface methodology we could enhance the production of antimycobacterial molecules chrysomycin A and B significantly.

**Importance:** Rediscovery of existing molecules and lack of techniques to induce production of secondary metabolites are the major bottlenecks associated with drug discovery of novel bioactive molecules from *Streptomyces*, the major source of marketable drugs today. We found a new method to increase the diversity and quantity of secondary metabolites in two *Streptomyces* species. This method thus enhances the chance of finding novel active principles from *Streptomyces*.

## 1. Introduction

From enzymes to antibiotics, most of the usable bio-active molecules have been discovered from Actinomycetes. However, there prevails a lameness in finding novel molecules from these organisms which led to the recent disinterest in this platform. This bottleneck is fuelled by the rediscovery of known molecules and lack of techniques in inducing the bacteria to produce novel products. Among them, *Streptomyces* are still an attractive source for the discovery of novel bio-actives. Their diverse biosynthetic gene clusters are responsible for the production of novel chemical entities (1). However, there seems to be a difficulty in inducing the expression of cryptic gene clusters coding for enzymes responsible for the production of diverse secondary metabolites under laboratory conditions. This failure in inducing these gene clusters has led to missing out potential secondary metabolite producers.

Conditioning morphology is a technique used to enhance secondary metabolite production (SMP) in *Streptomyces* by manipulating its growth conditions. For example, *Streptomyces* mycelia generally form aggregates in liquid media and inhibition of this clumping and achieving homogenous growth lead to the synthesis of novel compounds. This was conventionally performed by placing solenoids in the shake flask to disintegrate mycelial aggregates(2, 3). Similar results were achieved by manipulating genes such as *cslA, glxA* and *dtpA* to prevent aggregation without physical intervention. On the other hand, there are also contrary findings that show aggregation favors production of novel compounds (4-7). However, inside a bioreactor, pellet formation results in slower growth, and culture heterogeneity contributes to suppression of SMP. Sporulation in liquid media is another process that accompanies mycelialaggregation which also retards SMP. Thus, a conditioning method which disintegrates clumps, avoids sporulation and induces robust SMP is the need of the hour to rejuvenate the *Streptomyces* platform for finding novel bio-actives.

In this study, we a present a method for fermentation by *Streptomyces* which enhances and diversifies the SMP that eventually turns non-producers into producers of antibiotics. The method involves interference in cellular events of the life cycle of *Streptomyces* and which induces the SMP. We provide evidence for spatial separation of aerial mycelia from vegetative mycelia using glass marbles in the production flask. In addition, we observed inhibition of SMP by aerial mycelia, and that it is essential to remove them continuously from the fermentation medium to increase SMP. Earlier in our drug discovery program to discover antimycobacterial and anti-cancer molecules, we used this method and we could isolate two novel compounds (chrysomycin A & B - antimycobacterial molecules, and urdamycin – an anti-cancer agent). Using the method described here and response surface methodology we could scale up the production of chrysomycins by several fold compared to conventional methods of fermentation. Thus, the presence of marbles significantly increases the production of secondary metabolites.

## 2. Materials and Methods

### 2.1. Bacterial cultures and general materials

*Streptomyces sp.*OA161 and *Streptomyces sp.*, OA293 were from our own Actinomycetes repository (GenBank IDs KX364040 and KY014435, respectively). Polycarbonate autoclavable 250 mL Erlenmeyer flasks were procured from Tarsons(India) and used for culturing. Glass marbles, purchased locally, were 1 cm in diameter and weighed approximately 5 grams each. Nile red and SYTO 9 stains were procured from Sigma and Thermo Scientific, respectively. HPLC grade D-(+)-trehalose dehydrate, was procured from Fluka™ Fischer Scientific.

### 2.2. Fermentation

#### 2.2.1. Seed culture

The spores of *Streptomyces* isolates grown on solid ISP-2 (International Streptomyces Project medium) were used to prepare the seed culture. Seed cultures were grown in 50 ml of ISP-2 broth in 250mL flasks. Two sterile glass marbles were introduced into the flasks. The seed culture was incubated at 30 °C on a shaker incubator at 250 rpm for 72 h. The seed culture was used to inoculate 50 mL of fresh medium in test flasks at 1% (vol/vol) concentration.

### 2.3. Assessment of secondary metabolite profile

Fifty millilitre of ISP-2 medium was used as the fermentation medium for both the strains. Each 250 mL flask received 1% inoculum (vol/vol). Flask 1 contained none of the mechanical disruptors; flask 2 contained a solenoid immersed in the medium; and flask 3 and 4 contained two glass marbles in each. Generally, culturing with marbles resulted in the formation of a thick deposit on the inside of the flask due to the swirling motion of the medium during incubation. The deposit was pushed back into the medium in flask 4 using a sterile loop whenever it was formed. Flasks 5 and 6 were replicates of flask 1 until day 6. Later, flask 6 received the thick deposit from the wall of flask 3, and flask 5 served as control for this experiment. Both the flasks were incubated for 3 more days. After fermentation, 10 mL of media was drawn from each flask and the cell-free supernatant was extracted with equal volume of ethyl acetate by two-phase extraction. The ethyl acetate extract was collected and dried using a vacuum concentrator at 37°C. The concentrated extract was resuspended in 100µL of ethylacetate from which 10µL was spotted on to silica plates for performing TLC.

#### 2.3.1. Thin Layer Chromatography

Two-dimensional thin layer chromatography was performed to visualize and compare the SMP profile from the test and control flasks. Silica gel 60 F254 (Millipore India) was used as the stationary phase. In the first dimension, either hexane:acetone (1:1) or chloroform: acetone (7:3) was used for separating the compounds in the extracts from the culture filtrate of *Streptomyces sp.*, OA161 or *Streptomyces sp.*, OA293, respectively(8, 9). After drying at room temperature, the plates were turned 90 degrees, and ethyl acetate was used to resolve molecules in the second dimension. After drying, photographs were taken using Light L16 camera under UV light (254 nm). Densitometry analysis of the spotswas performed with Image J software. The results were plotted and tested for significance using GraphPad Prism version 7.

#### 2.3.2. LC- MS analysis

##### 2.3.2.1. Analysis of metabolites in ethylacetate extracts

The concentrated extracts prepared for TLC were resuspended in 400µL of acetonitrile instead of ethyl acetate. Each sample (7.5 µL) was injected into the C18 reverse phase (high-strength silica, 2.1 × 100 mm, 1.8 µm) Waters column maintained at 40 °C. The mobile phase had two components - 0.1% formic acid in ultrapure water (A) and 0.1% formic acid in acetonitrile (B). The run program followed was 0 min, 1% B; 2 min, 10% B; 6 min, 30% B; 8 min, 50% B; 12 min, 75% B; 15 min, 99% B; and 20 min, 1% B. Further analysis was performed using Waters ACQUITY UPLC system (Waters, Milford, MA, USA) coupled to a quadrupole–time-of-flight (Q–TOF) mass spectrometer (SYNAPT-G2, Waters). The total run was for 20 min and the systems were operated and controlled by MassLynx4.1 SCN781 software (Waters).

##### 2.3.2.2. Mass spectrometry

The SYNAPT® G2 High Definition MS™ System mass spectrometer (Waters) was used in positive and negative resolution mode with electrospray ionization (ES+) source over a mass range of 50–1500 Da. The capillary voltage and sampling cone voltage were set at 3.0 kV and 30 kV, respectively. Cone and desolvation gas flow rates were adjusted to 80 L/h and 600 L/h, respectively. Ion source and desolvation gas (nitrogen) temperatures were kept constant at 130 °C and 450 °C, respectively. Lock mass acquisition was performed every 30 s by leucine– enkephalin (556.2771 [M+H]+) for accurate on-line mass calibration. All the spectra were acquired for 0.25 s with an inter-scan delay of 0.024 s. The total metabolite profiles were compared for differential production of compounds. In addition, random molecular weights were compared to see if there is a differential production in specific molecules.

### 2.4. Quantification of trehalose content

The thick deposits on the inside wall of the flasks and the broth culture (wet weight of 150 mg of each) were washed with 1X PBS (phosphate buffered saline). After washing, sterile water (1 mL) was added to the bacterial pellet and boiled for 30 min in a boiling water bath. The suspension was passed through 0.2 micron filter and the contents were separated using HILIC column. Acetonitrile:ammonium acetate gradient was used as the mobile phase and the total duration of the run was 22 minutes. MS was employed to identify trehalose, HPLC-grade trehalose, Sigma served as the standard.

### 2.5. Nile Red Staining and Confocal Microscopy

Nile red staining was performed as described elsewhere(10)on the broth culture and the bacteria from the circular depositof both the flasks (with and without glass marbles). Briefly, bacterial smear on a glass slide was flooded with Nile red dissolved in methanol (40 µg/mL). After 20 minutes, it was washed with running water and counterstained with SYTO 9. The slides were visualized under a Nikon-A1 R confocal microscope.

### 2.6. Scanning electron microscopy

*Streptomyces* culture from the circular deposit from flask 3 was loaded on to gelatin coated cover-slips and then fixed overnight with 6.25% glutaraldehyde (in 50 mM phosphate buffer, pH 7.4). The cover slips were then dehydrated using increasing gradients of ethanol (30-100%) and left for drying. The dried cover slips were then fixed on stubs and sputter-coated with gold followed by imaging using Scanning electron microscope (FEI ESEM Quanta 200-3D, USA).

### 2.7. Analysis of cell death

Cultures incubated for 4-6 days were subjected to cell death analysis. LIVE/DEAD®BacLight™ bacterial viability kit procured from Invitrogen (L7007 Molecular Probes, Invitrogen) was used to assess cell death in the strains grown by different methods. Procedures were performed as per the instructions of the manufacturer. Briefly, the bacterial cells were incubated with reagent mixture of the kit at 37°C for 15 min in the dark and the fluorescence intensity was measured immediately using TECAN Infinite M200 (data acquired by Magellen v6.6 software) at 530 nm (green) and 630 nm (red), after excitation at 485nm. The ratio of red to green intensity was calculated and compared for the cultures grown with and without marbles.

### 2.8. Media optimization

#### 2.8.1. Bacterial culture conditions and estimation of chrysomycin

Initially, *Streptomyces sp*.,OA161 - seed culture was prepared by inoculating a loop-full of spores into ISP-2 medium.After incubation for 72 h, 1% (v/v) inoculum was used to grow the bacteria on media containing different carbon sources (ISP-1, ISP-2, mISP-2, ISP-4, and ISP-7). The cultures were incubated at 28°C for 6 days at 250 rpm on a shaker incubator. After fermentation, the bacteria-free medium was extracted with equal volume of ethyl acetate and concentrated. The antimycobacterial fraction (AMF) was separated as described in section 2.3.1 (9). Intrinsic fluorescence of chrysomycin A & B, was used for the assessment of their production. Upon shining UV light (365 nm), the green fluorescence emitted by chrysomycins was photographed and densitometry analysis was performed with Image J software. The results were plotted and tested for significance using GraphPad Prism version 7.

#### 2.8.2. Screening for essential media components for production of chrysomycin A & B by Plackett-Burman Design (PBD)

PBD was employed for identifying the significant factors that influence chrysomycin production. To increase the robustness of the design, a three level factorial design with high (+1), intermediate (0) and low (−1) levels with 4 centre points was used in the screen compared to the conventional two factorial design. The matrix design is shown in Table 1, and the media components along with their concentrations are provided in Table 2. The influencing factors were screened by performing an F-test. A Pareto chart was also generated with the t-value of the effect to representhow each factor affects the production of chrysomycin. The positive influencing factors were used in the interaction study with Box-Behnken designs. Negatively influencing factorsand factors that induced dummy variable traps were not considered forsubsequent studies. Components of modified ISP-2 medium, sodium chloride and calcium carbonate along with dummy factors (magnesium sulphate, ferrous sulphate, asparagine, sodium hydrogen phosphate, Tween 80 and temperature) were selected for the screening process. All the experiments were conducted in triplicates.

**Table 1.**
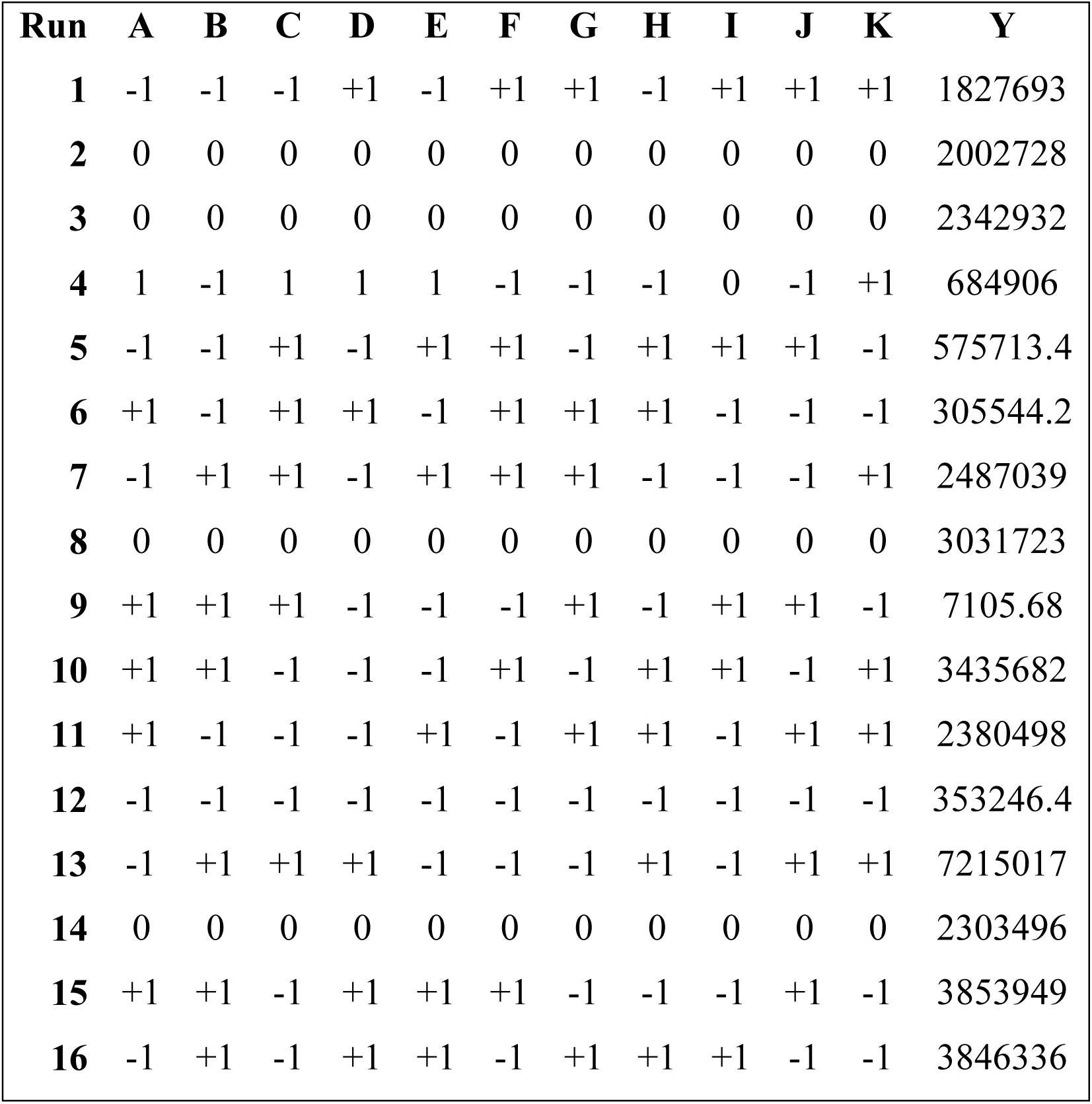
Plackett-Burman matrix of the experimental design. A to K represent test factors and Y denotes the response. A: maltose; B: yeast extract; C: dextrose; D: calcium carbonate; E:temperature; F: sodium chloride; G: magnesium sulphate; H: ferrous sulphate; J: Tween 80; K: asparagine; L:sodium dihydrogen phosphate; Y: chrysomycins production.

| **Run** | **A** | **B** | **C** | **D** | **E** | **F** | **G** | **H** | **I** | **J** | **K** | **Y** |
| --- | --- | --- | --- | --- | --- | --- | --- | --- | --- | --- | --- | --- |
| **1** | -1 | -1 | -1 | +1 | -1 | +1 | +1 | -1 | +1 | +1 | +1 | 1827693 |
| **2** | 0 | 0 | 0 | 0 | 0 | 0 | 0 | 0 | 0 | 0 | 0 | 2002728 |
| **3** | 0 | 0 | 0 | 0 | 0 | 0 | 0 | 0 | 0 | 0 | 0 | 2342932 |
| **4** | 1 | -1 | 1 | 1 | 1 | -1 | -1 | -1 | 0 | -1 | +1 | 684906 |
| **5** | -1 | -1 | +1 | -1 | +1 | +1 | -1 | +1 | +1 | +1 | -1 | 575713.4 |
| **6** | +1 | -1 | +1 | +1 | -1 | +1 | +1 | +1 | -1 | -1 | -1 | 305544.2 |
| **7** | -1 | +1 | +1 | -1 | +1 | +1 | +1 | -1 | -1 | -1 | +1 | 2487039 |
| **8** | 0 | 0 | 0 | 0 | 0 | 0 | 0 | 0 | 0 | 0 | 0 | 3031723 |
| **9** | +1 | +1 | +1 | -1 | -1 | -1 | +1 | -1 | +1 | +1 | -1 | 7105.68 |
| **10** | +1 | +1 | -1 | -1 | -1 | +1 | -1 | +1 | +1 | -1 | +1 | 3435682 |
| **11** | +1 | -1 | -1 | -1 | +1 | -1 | +1 | +1 | -1 | +1 | +1 | 2380498 |
| **12** | -1 | -1 | -1 | -1 | -1 | -1 | -1 | -1 | -1 | -1 | -1 | 353246.4 |
| **13** | -1 | +1 | +1 | +1 | -1 | -1 | -1 | +1 | -1 | +1 | +1 | 7215017 |
| **14** | 0 | 0 | 0 | 0 | 0 | 0 | 0 | 0 | 0 | 0 | 0 | 2303496 |
| **15** | +1 | +1 | -1 | +1 | +1 | +1 | -1 | -1 | -1 | +1 | -1 | 3853949 |
| **16** | -1 | +1 | -1 | +1 | +1 | -1 | +1 | +1 | +1 | -1 | -1 | 3846336 |

**Table 2.**
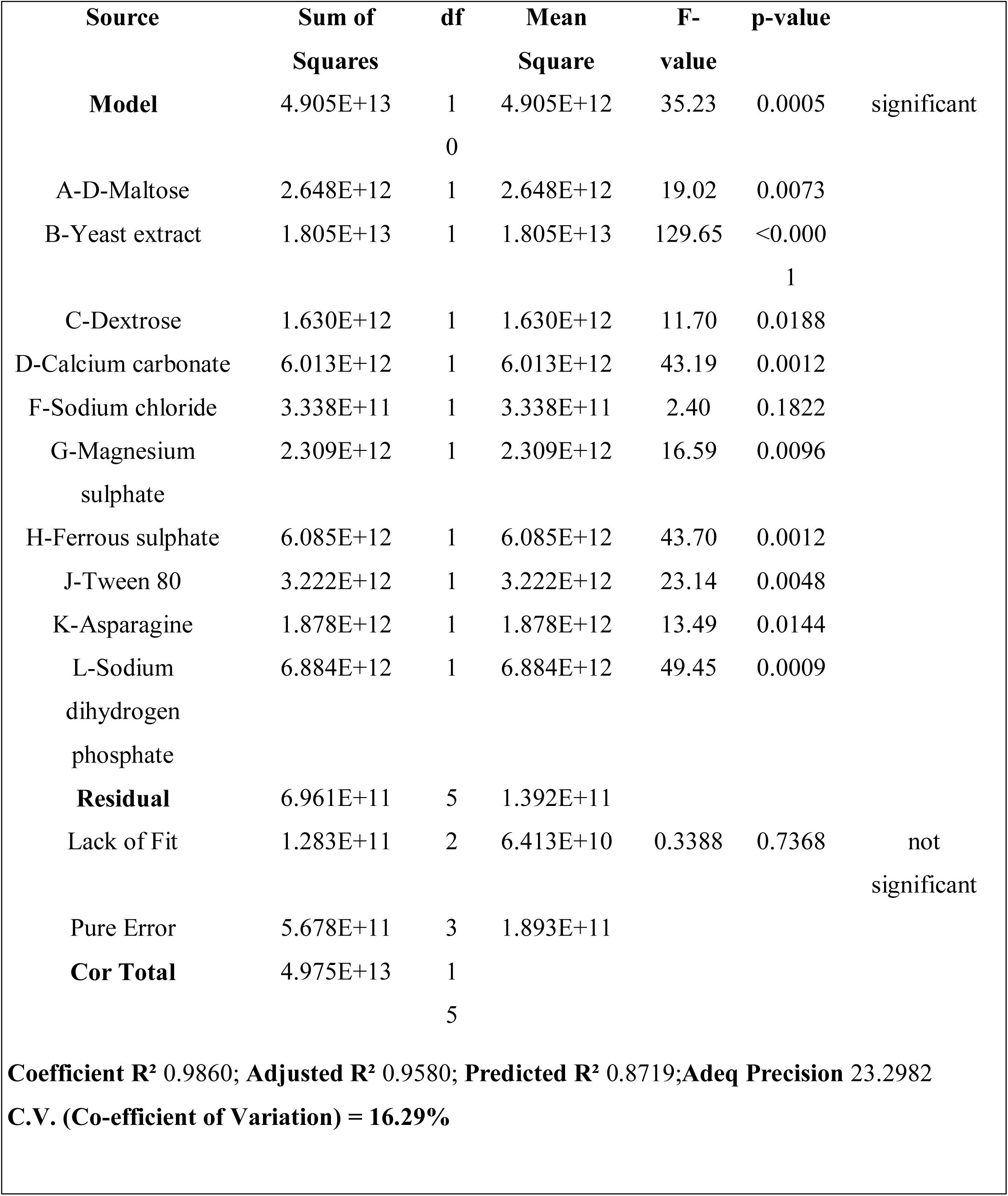
Results of ANOVA for Plackett-Burman designs.

#### 2.8.3. Optimization of medium components by Response Surface Methodology

Box-Behnken design was used to study the interaction between the screened factors with respect to chrysomycin production. A three-level factorial design with 2 centre points was used in the interaction study. The data were analyzed by fitting the regression values into a second order polynomial equation,

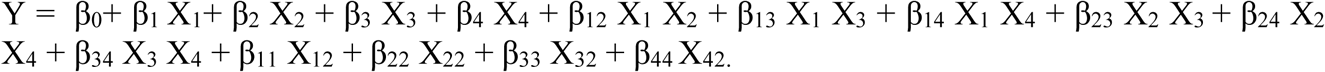

Y represents the response values, β denotes the regression coefficient, and X_1_, X_2_, X_3_, X_4_ represent the significant factors selected in this study. The character of the polynomial fit is expressed as coefficient R^2^, and the significance was calculated by F-test. All the experiments were conducted in triplicates. The design matrix containing the screened components and their respective concentrations are given in the electronic version of the supplementary data.

#### 2.8.4. Statistical Analysis

Design-Expert software 12.0 (*Stat-Ease, Minneapolis*) was employed for the experimental design and data analysis. Models were used to predict best yield of chrysomycin and the derived combinations of medium components were used to validate the effect of our observations on the production of chrysomycin A & B. The predicted values were compared with the observed values, and significance of the data was obtained to validate the models.

## 3. Results and Discussion

### 3.1. Marbles assist in robust SMP in*Streptomyces sp*

Figure 1A shows schematic representation of flasks with and without mechanical disruptors. Figures 1B and 1C represent 2D TLC profiles of secondary metabolites of bacteria-free supernatants of two different isolates (*Streptomyces* sp., OA161 and *Streptomyces* sp., OA293). Medium in flask 3 (with marbles) had more secondary metabolites than flask 1 and 2, and was supported by LC-MS analysis. In addition, comparision of profiles showed that production of the metabolites was enhanced and production of a few metabolites remained unaffected (Supplementary Figure 1). We speculate that the latter are basal metabolites involved in bacterial survival. The strains used in the experiments were known to produce unique antibiotics, chrysomycins (9) and urdamycins, respectively (11). Therefore, we compared their production by 2D TLC (white arrows point at the compounds, Figures 1D and 1E) to support the enhanced SMP. As expected these compounds were produced in very small quantities or absent in conventionally grown cultures as against their significantly high levels when marbles were introduced into the cultures.

**Figure 1:**
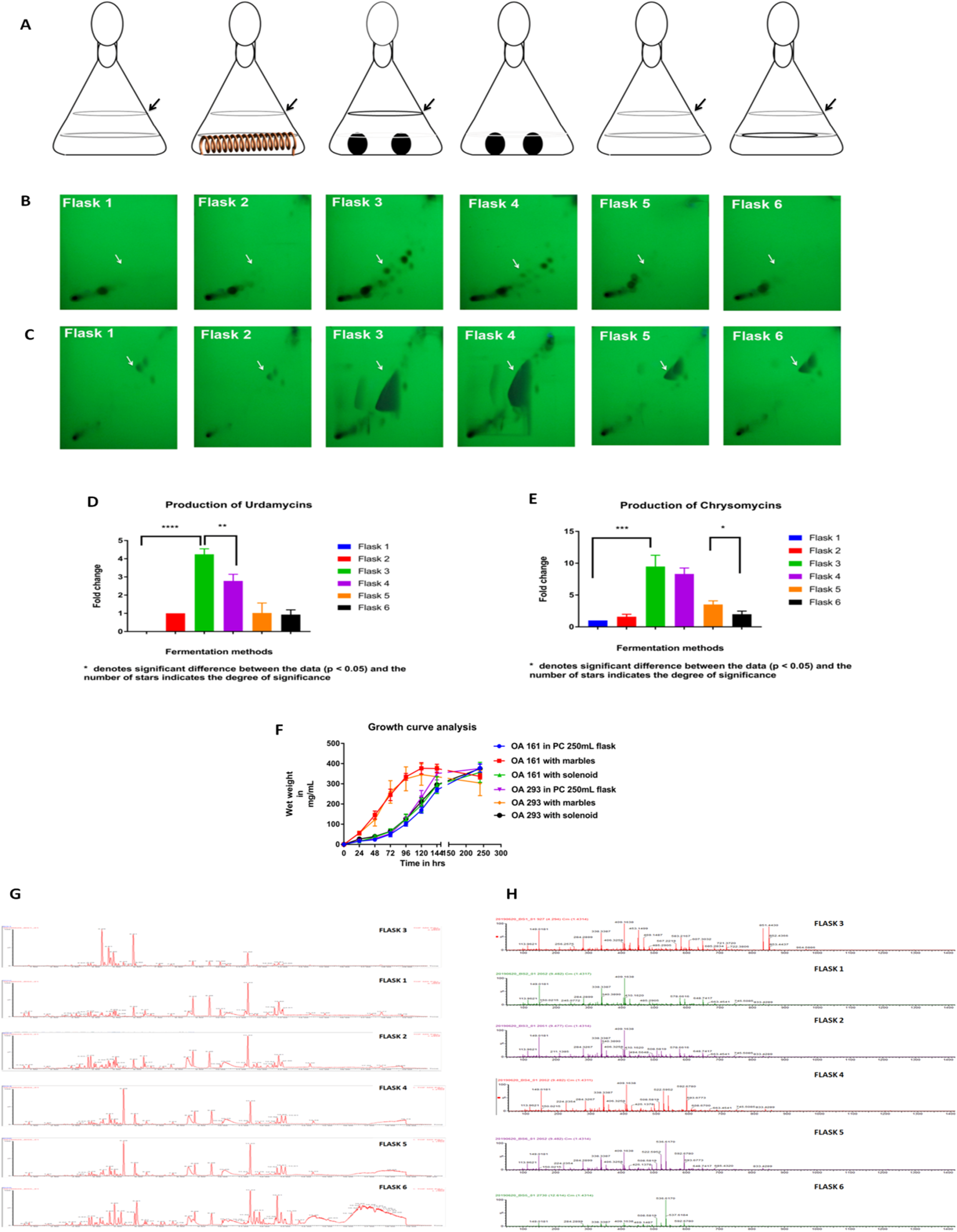
Marbles assist in robust SMP by separating the production-retarding mycelia from the metabolically active mycelia. Schematic representation of the test shake flask cultures is given in A. B and C represent secondary metabolite profiles (2D TLC) from *Streptomyces* sp.OA161 and *Streptomyces* sp.,OA293, grown in flasks as represented above. White arrow denotes urdamycins in B, and chrysomycins in C. D and E represent the densitometric measurement of B and C, respectively. F shows thegrowth curves of both the *Streptomyces* species grown with and without marbles. G and H show the HPLC and mass spectrometry profiles of the ethyl acetate extracts of flasks 1 to 6.

### 3.2. Marbles increase the growth rate of*Streptomyces* and assist in spatial separation of bacterial mycelia

To analyze whether marbles provide an advantage in biomass production, the growth pattern of both the isolates grown in flasks 1, 2 and 3 were tracked and compared (Figure1E). Interestingly, the strains had insignificant difference in biomass, but differed in their growth rates. Generally a biphasic growth is commonly observed in most *Streptomyces* cultures (4). However, in the presence of marbles, cultures exhibited a standard sigmoid growth curve. This was intriguing and warranted the examining of the morphology of the culture under different growth conditions. Interestingly, all the flasks had bacterial deposit on their inside as shown in the representative figure (Figure 1A), but flask 3 had the thickest ring and had a spore-like chalky appearance.

### 3.3. The bacterial deposits suppress SMP on contact withthe broth culture

To find whether the bacterial deposit on the wall (flask 3) affect SMP, the ring was pushed back into the broth culture whenever visible aggregates were formed (until 6 days). Interestingly, this significantly reduced the SMP (flask 4 in Figures 1B and 1C). To confirm that this reduction was indeed because of the bacterial deposit, these were transferred to a flask without any mechanical disruptors (Flask 6) which was previously incubated for 6 days. As expected, a similar reduction pattern was observed in flask 6 compared to flask 5. This data was in line with the data shown in flask 3 and 4 (Figure 1E).The metabolomic patterns observed in 2D TLC plate were validated using LC-MS analysis (Figure 1F and 1G) and the number of metabolites were compared and presented in supplementary figure 2.

### 3.4. The deposit on the inside wall of the flask was identified as the aerial mycelium of *Streptomyces*

In the life cycle of *Strepomyces*, a spore germinates to form vegetative mycelium during favorable conditions and later differentiates into aerial mycelium. Progression of spore formation in aerial mycelia is accompanied by formation of a hydrophobic sheath around them during which SMP diminishes. Hence, we hypothesized that the marbles could be interfering with this step to enhance SMP. They could be aiding in selectively removing the aerial mycelium from the vegetative mycelia. To test this possibility, we stained the aerial and vegetative mycelia with Nile Red (which is used to stain hydrophobic and neutral lipids) to distinguish between hydrophilic and hydrophobic mycelia. In Figure 2A, the first two rows represent *Streptomyces* sp.,OA161 from broth culture, and the rest represent bacteria from the wall deposits. The green color shows the presence of the hydrophilic bacteria (as SYTO9 stains the DNA) and red represents the hydrophobic bacteria stained by Nile red. The broth culture from flask 1 had intense red stain throughout while culture of flask 3 had picked up the red stain in isolated regions. On comparing the bacteria from wall deposits, red stain was absent on the thin wall deposits of flask 1 while the thicker counterpart of flask 3 stained intense red. Similar results were obtained when the same experiment was repeated on *Streptomyces* sp.,OA293 (Supplementary Figure 3). These results suggest that the marbles caused mechanical shearing that separated the aerial mycelium. The hydrophobic aerial mycelia were deposited on the sides of the flask due to the swirling movement of the medium. On the other hand, in flasks without marbles, vegetative mycelium was found forming insignificant amounts of deposits which did not affect the SMP. Further to confirm whether the intense red region is in fact the broken aerial mycelium, scanning electron microscopy was performed which revealed aerial mycelium with spores (Figure 2B). Manipulation of genes such as *csl* A and *glx* A to prevent differentiation of aerial mycelium was attempted recently (12) to increase SMP. Marbles seem to provide a simple and inexpensive alternative to such genetic manipulations and provide better yield of secondary metabolites.

**Figure 2:**
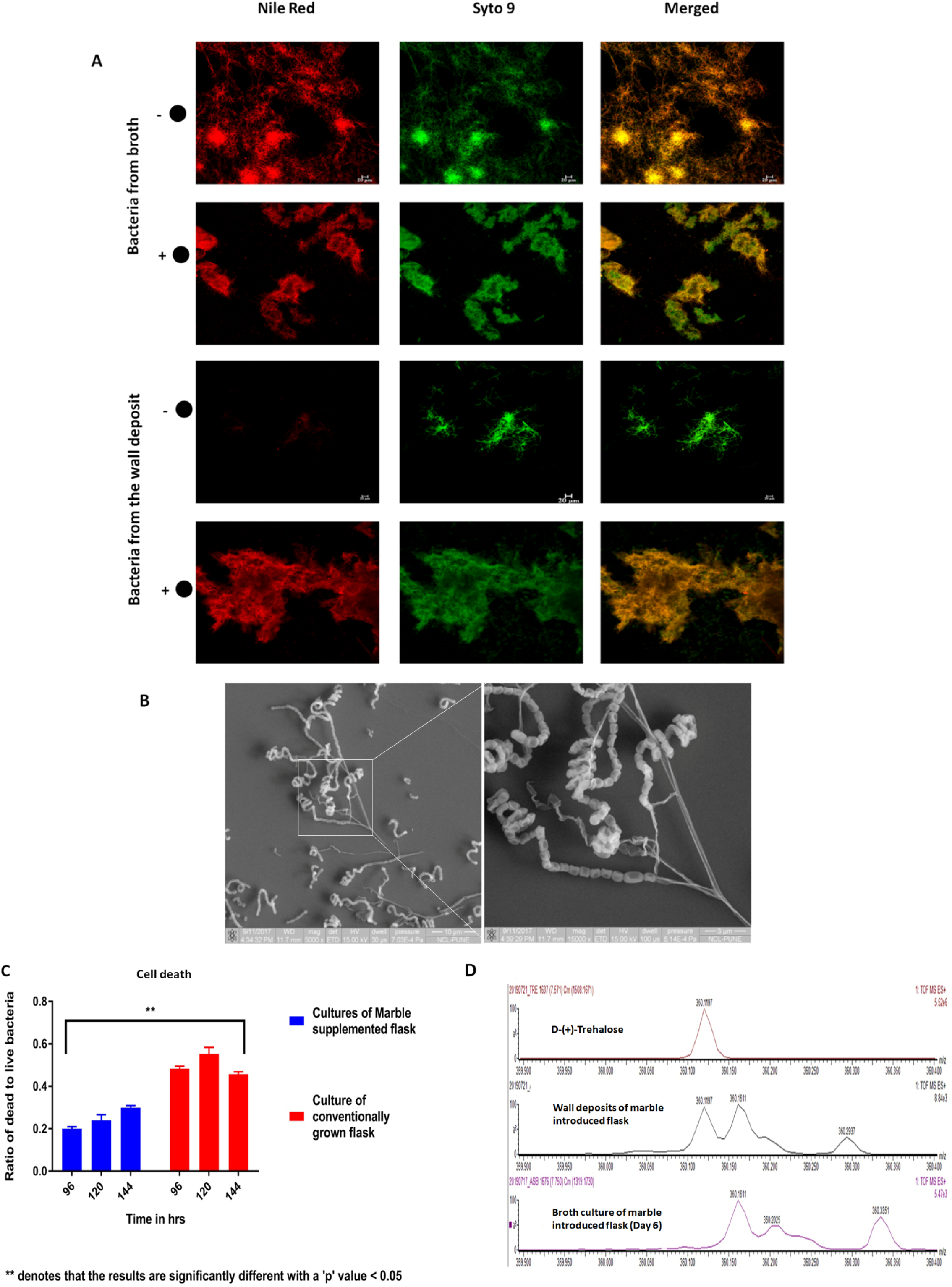
The wall deposits areaggregates of aerial mycelium. (A) Confocal microscopy images show bacteria from broth and wall deposits. ‘+●’ and ‘-●’ represent bacteria from flasks with and without marbles, respectively. The differentiated aerial mycelium appears red and all live bacteria are green. (B) Scanning electron micrographs of the deposit on the inner wall of the flask. The expanded image on the right shows the morphology of the sporophores on the aerial mycelia. (C) Graphical representation of cell death analysis. (D) Comparison of trehalose content of wall deposits and the broth culture, by LC-MS analysis.

### 3.5. Measurement of cell death and intracellular trehalose

In the life cycle of *Strepomyces*, after the initiation of aerial mycelium formation, the vegetative mycelium undergoes cell death. Therefore, cell death was assessed inbroth cultures of flasks 1 and 3 during 96 h to 144h (stationary phase). The ratios of dead to live bacteria were lower in the presence of marble (flask 3) supporting better SMP. On the other hand, the ratios were significantly higher in flask 1 correlating with the formation of aerial mycelium (Figure 2C). Trehalose content in the mycelium is known to reflect the state of development of *Streptomyces*. Increased trehalose content is reported in differentiated aerial mycelium compared to glycogen-rich vegetative mycelium (13, 14). Therefore, we compared the trehalose content of the broth culture and wall deposits of flask 1 and 3. As expected, the trehalose content of the wall deposit was higher than that in the broth cultures of flask 3 (Figure 2D, and Supplemetary Figure 3A). However, we could not establish the same in the other strain, *Streptomyces* sp., OA293, as different *Stretomyces* species are known to use different storage molecules. For example, *S. lividans* was found to accumulate triacylglycerols instead of trehalose (15). Thus, the results of cell death and trehalose accumulation support the hypothesis that the marbles selectively remove the differentiating aerial mycelia and thereby extend the period of SMP. Although this study found a robust method for SMP, we could not establish a link between the enhanced SMP and activation of cryptic biosynthetic gene clusters in these organisms.We assume that the stress caused by breakage of mycelia might have triggered the induction of cryptic genes leading to enhanced SMP. Further, the results call for the designing of non-canonical bioreactors which can selectively remove the aerial mycelium from the broth culture to increase the yield. Based on our observations, we propose that by incorporating novel designs in fermentors to remove aerial mycelia from *Streptomyces*, it is possible to enhance the yield of secondary metabolites while scaling up.

### 3.6. Optimization of chrysomycin production medium through response surface methodology

#### 3.6.1. Selection of media and conditions for antibiotic production

The production of chrysomycins was better in ISP-2 medium compared to that in other media (ISP-1, ISP-4 and ISP-7) tested. However, when malt extract was replaced with maltose in ISP-2 medium (named as modified ISP-2 medium (mISP-2)), the production of chrysomycins was found to increase significantly (Figure 3A). This encouraged us to use the components of the modified medium in response surface methodology (RSM) studies. One-factor-at-a-time approach ensured that variation in temperature had least effect on production (Figure 3B), and thus it was included as a dummy factor/variable. Altogether, the main components of the mISP-2 medium (maltose, yeast extract and dextrose) and dummy factors such as Tween80, sodium hydrogen phosphate, temperature, magnesium sulphate, ferrous sulphate and asparagine were included in the study. As the *Streptomyces* OA161 strain was mildly halophilic (Supplementary Figure 4C) we included sodium chloride to the list of test variables. Calcium carbonate was also included in the study as it is known to enhance SMP in actinomycetes (16).

**Figure 3:**
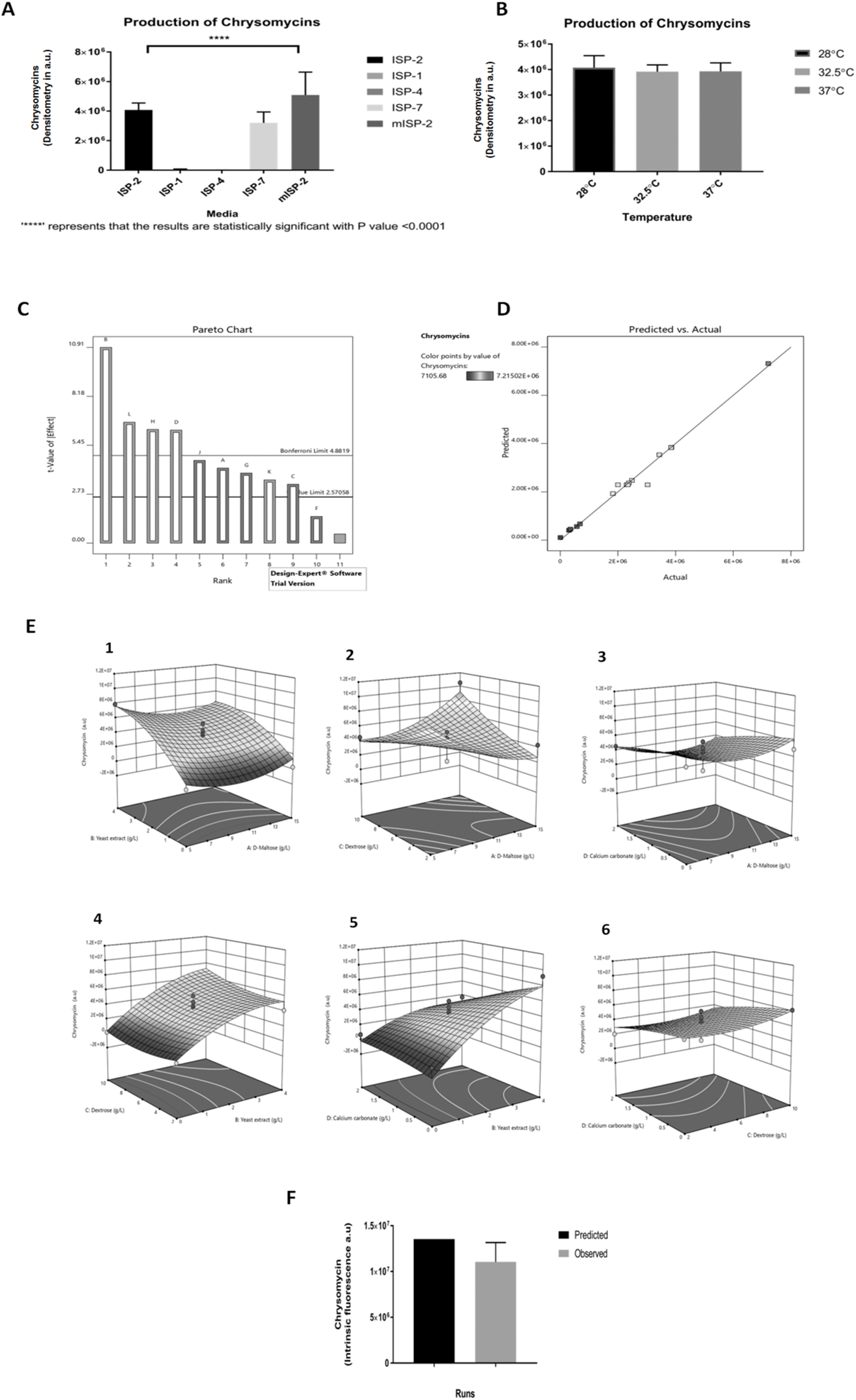
Optimization of chrysomycin production media. (A) Selection of basal medium for the production of chrysomycin. (B) Effect of temperature on production of chrysomycin. (C) Pareto chart showing the effect of individual variables on chrysomycin production. (D)Diagnostic plot of the Plackett-Burman design model shows that predicted and observed responses fall on a straight line. (E) Three-Dimensional plots showing interaction between medium components on chrysomycin production. 1. yeast extract vs maltose; 2. dextrose vs maltose; 3. calcium carbonate vs maltose; 4. dextrose vs yeast extract; 5. calcium carbonate vs yeast extract; 6. calcium carbonate vs dextrose. (F)The predicted and observed levels of chrysomycin production after optimization.

#### 3.6.2. Screening of factors that influence production of chrysomycins

Plackett-Burman design (PBD) was used for the screening process. A total of 16 experiments were performed as suggested by the PBD for screening 11 variables. Their individual responses (chrysomycins production in arbitrary units) are given in the Supplementary Table 1. The amount of chrysomycins was determined from the TLC plate (Supplementary Figure 4B). The relationship between the variable factors and responses were analyzed by analysis of variance (ANOVA). The statistical significance of the model and the variables are given in Table 2. Factors other than temperature and sodium chloride affected the production significantly. Generally, the alterations caused by the dummy factor in production and their positive statistical significance are results of ‘dummy variable traps’ and hence those factors were not consideredin subsequent studies. A Pareto chart (Figure 3C) was generated based on the t-values calculated and it showed that yeast extract and calcium carbonate enhanced production while the carbon sources dextrose and maltose exerted a negative influence on the production of chrysomycins. The predicted responses were in agreement with the observed responses as evidenced by a straight line in Figure 3D. This strongly suggests the need of an interaction study to formulate an efficient production media as dextrose and maltose are the major carbon sources. Regression values were calculated and fit into a linear equation as follows,

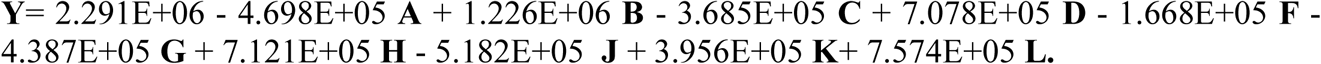

Furthermore, R^2^ value of 0.986 states that the model equation could justify 98.6% of the total variation.

#### 3.63. Optimization of medium components with Box-Behnken Designs (BBD)

As maltose, dextrose, calcium carbonate and yeast extract were identified as the most influential factors, they were used for optimizing the production medium (data provided in the Supplementary Table 2) using BBD. Twenty runs were performed at 30°C and 250 rpm. The effect of individual variables on production was assessed using ANOVA (Table 3). Briefly, the model was statistically significant and R^2^ value of 0.93 indicates that the model equation could justify 93% of the total variation. Also, the regression value for each variable was fitted into a second-order quadratic equation as a function of response values.

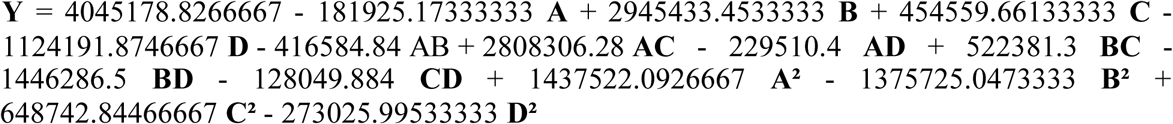

**Table 3.** Results of ANOVA for Box-Behnken designs.

| Source | Sum of Squares | df | Mean Square | F-value | p-value |  |
| --- | --- | --- | --- | --- | --- | --- |
| Block | 1.280E+13 | 2 | 6.402E+12 |  |  |  |
| **Model** | 1.996E+14 | 14 | 1.426E+13 | 12.34 | < 0.0001 | significant |
| A-D-Maltose | 3.972E+11 | 1 | 3.972E+11 | 0.3437 | 0.5678 |  |
| B-Yeast extract | 1.041E+14 | 1 | 1.041E+14 | 90.09 | < 0.0001 |  |
| C-Dextrose | 2.479E+12 | 1 | 2.479E+12 | 2.15 | 0.1667 |  |
| D-Calcium carbonate | 1.517E+13 | 1 | 1.517E+13 | 13.12 | 0.0031 |  |
| AB | 6.942E+11 | 1 | 6.942E+11 | 0.6007 | 0.4522 |  |
| AC | 3.155E+13 | 1 | 3.155E+13 | 27.30 | 0.0002 |  |
| AD | 2.107E+11 | 1 | 2.107E+11 | 0.1823 | 0.6764 |  |
| BC | 1.092E+12 | 1 | 1.092E+12 | 0.9446 | 0.3488 |  |
| BD | 8.367E+12 | 1 | 8.367E+12 | 7.24 | 0.0185 |  |
| CD | 6.559E+10 | 1 | 6.559E+10 | 0.0568 | 0.8154 |  |
| A<sup>2</sup> | 1.417E+13 | 1 | 1.417E+13 | 12.26 | 0.0039 |  |
| B<sup>2</sup> | 1.298E+13 | 1 | 1.298E+13 | 11.23 | 0.0052 |  |
| C<sup>2</sup> | 2.886E+12 | 1 | 2.886E+12 | 2.50 | 0.1381 |  |
| D<sup>2</sup> | 5.112E+11 | 1 | 5.112E+11 | 0.4423 | 0.5176 |  |
| **Residual** | 1.502E+13 | 13 | 1.156E+12 |  |  |  |
| Lack of Fit | 6.779E+12 | 10 | 6.779E+11 | 0.2467 | 0.9600 | not significant |
| Pure Error | 8.244E+12 | 3 | 2.748E+12 |  |  |  |
| **Cor Total** | 2.274E+14 | 29 |  |  |  |  |
| **Coefficient R<sup>2</sup> 0.9300; Adjusted R<sup>2</sup> 0.8546; Predicted R<sup>2</sup> 0.7269; Adeq Precision 12.7578; C.V. (Co-efficient of Variation) = 25.47** |  |  |  |  |  |  |

Y is the response (chrysomycin A & B production) and A, B, C, D are factors denoting maltose, yeast extract, dextrose and calcium carbonate. The predicted responses were in agreement with the observed responses, indicated by a straight line. The response surface plots were generated as three-dimensional contour plots illustrating the interaction between two factors individually keeping the other factors at their mid-level. The diagnostic plots that are indicators for the validated model is shown in Supplementary Figure 4D. Analyzing the 3D plots, there seemed to be no interaction between calcium carbonate and dextrose; calcium carbonate and yeast extract; and between maltose and yeast extract, respectively (Figure 3E). However, a positive interaction was observed between maltose and dextrose at the low and high concentrations but not at the mid level concentration. This is in line with our earlier observation that when malt extract is replaced by maltose, it leads to higher production of chrysomycins. Similar interaction was also observed between dextrose and yeast extract. Also, a negative interaction was observed with calcium carbonate and maltose. Considering the results, a list of solutions were sought from the Design Expert software which suggested 15 mg/mL of maltose, 4 mg/mL of yeast extract and 10 mg/mL of dextrose with no calcium carbonate in the production media can lead to a yield of 1.35429E+07a.u. of chrysomycins. The same was observed in fermentation (Figure 3F) when the predicted and observed responses were compared. Also, no significant difference was observed between them. Thus, the model was validated and a three-fold increase in production was achieved. This is the first report that attempted to optimize production of chrysomycins through statistical methods. We could successfully enhance the production to almost 24 fold when compared to traditional fermentation techniques.

## 4. Conclusion

Introduction of glass marbles in culture flasks results in robust secondary metabolite production in *Streptomyces*. The marbles spatially separate the differentiated aerial mycelia from vegetative mycelia by mechanical shearing and the swirling movement of the medium physically lifts the aerial mycelium along the walls of the flask. This method could be used for effective SMP in *Streptomyces*. We speculate that this separation of aerial mycelium can lead to the activation of biosynthetic gene clusters responsible for SMP. Response surface methodology helped in optimizing the media components for increased yield of chrysomycins. We achieved an overall 24 fold increase in production when compared to the conventional methods of fermentation.

## Author contributions

RAK and BM conceptualized and wrote the manuscript. BM, RRJ, RR, NM and LP performed the experiments.VJS and SD performed SEM analysis. KBA and VMD made the tables and figures.

## Funding

## Acknowledgement

BM thanks CSIR, Govt. of India for research fellowship and RAK thanks Department of Biotechnology, Govt. of India for funding. The authors thank Dr. Rajeev K. Sukumaran, Microbial processes and technology division, National institute for interdisciplinary Science and Technology (NIIST), Trivandrum for his critical evaluation of the manuscript.

**Supplementary data is attached as a separate word file**

## Conflict of interests

The authors declare no conflict of interests.

## Supplementary data

**Supplementary Figure 1:**
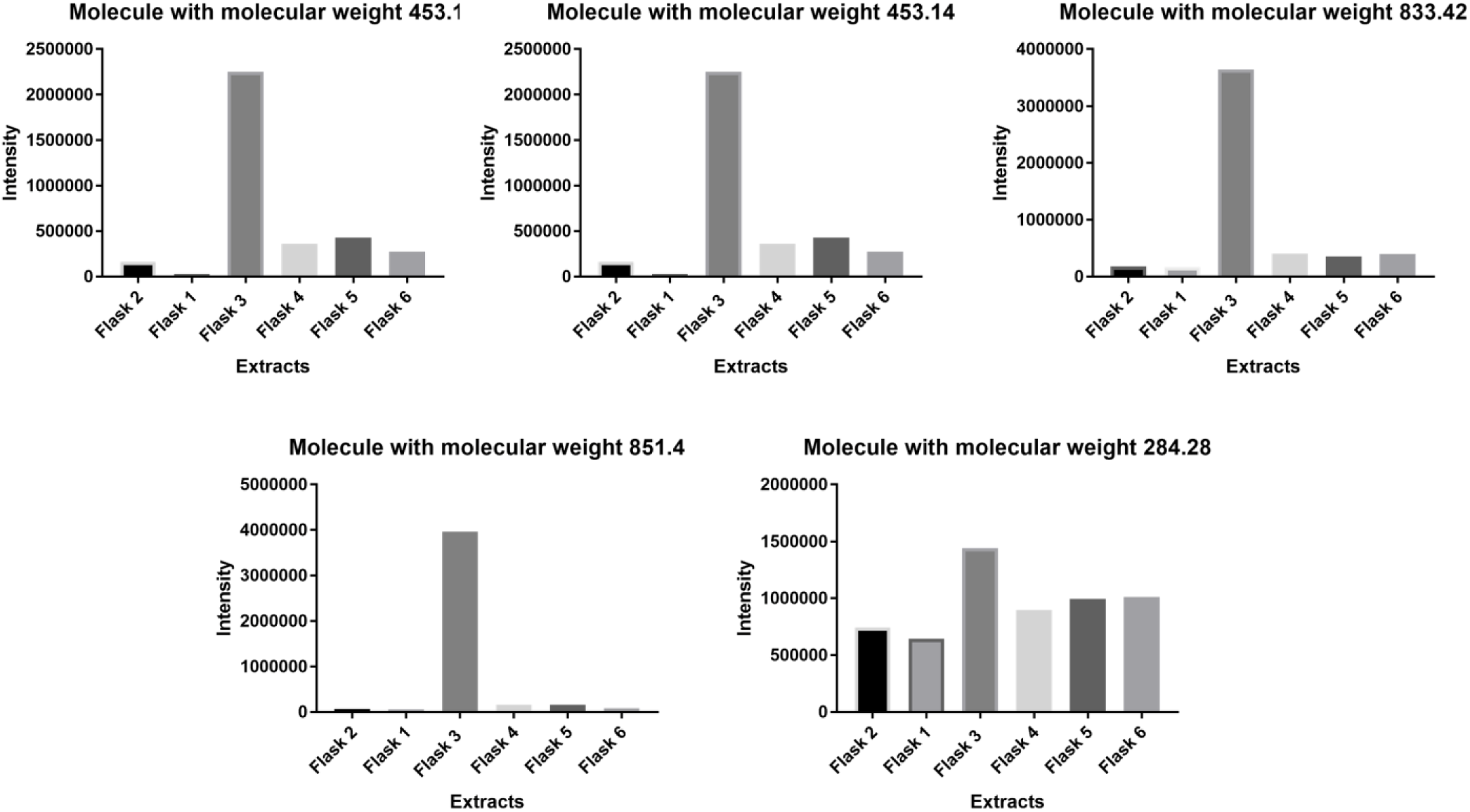
Representative images showing amount of random molecules present in extracts from different flasks. Molecule with molecular weight 284.28 had insignificant variations in amount while others follow similar pattern found in case of chrysomycins and urdamycins.

**Supplementary figure 2:**
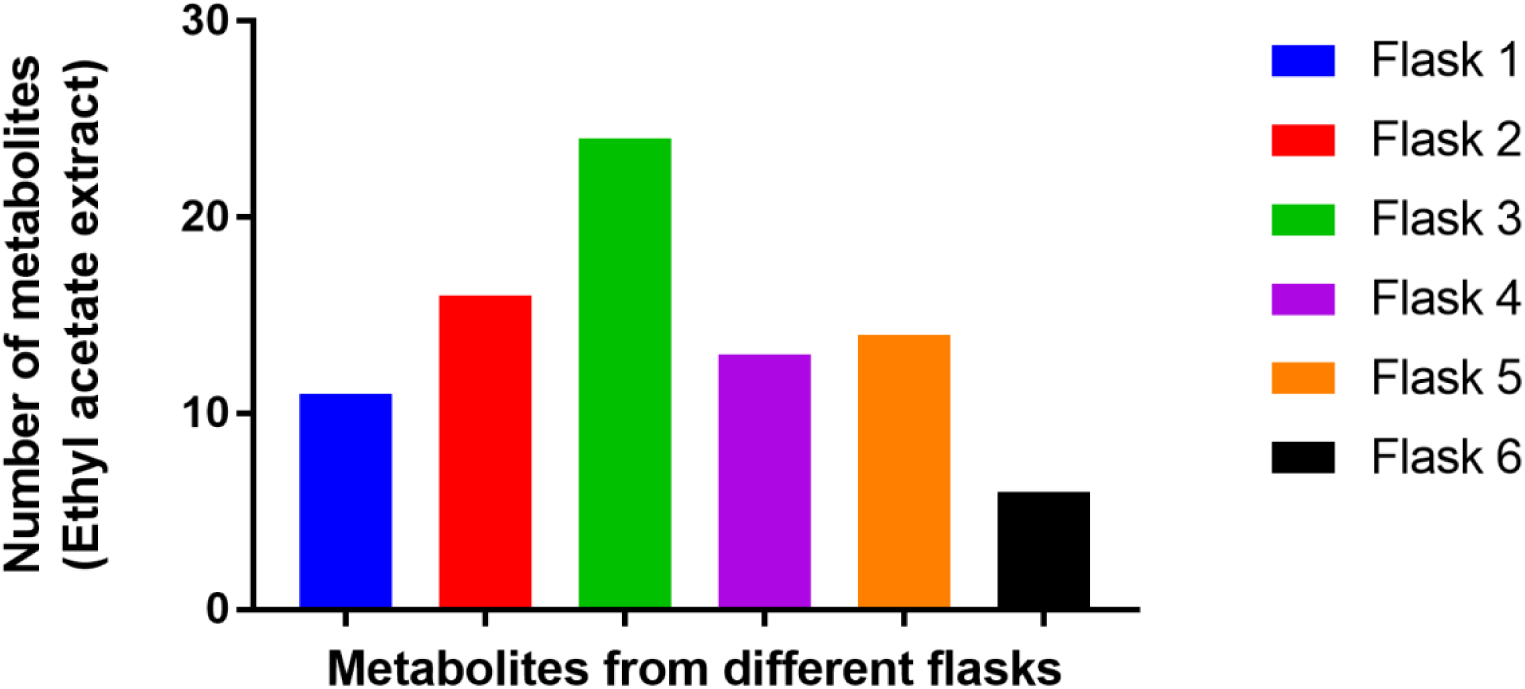
Difference in qualitative production of secondary metabolites of Ethyl acetate extracts of *Streptomyces* sp., OA161 culture filtrate.

**Supplementary Figure 3:**
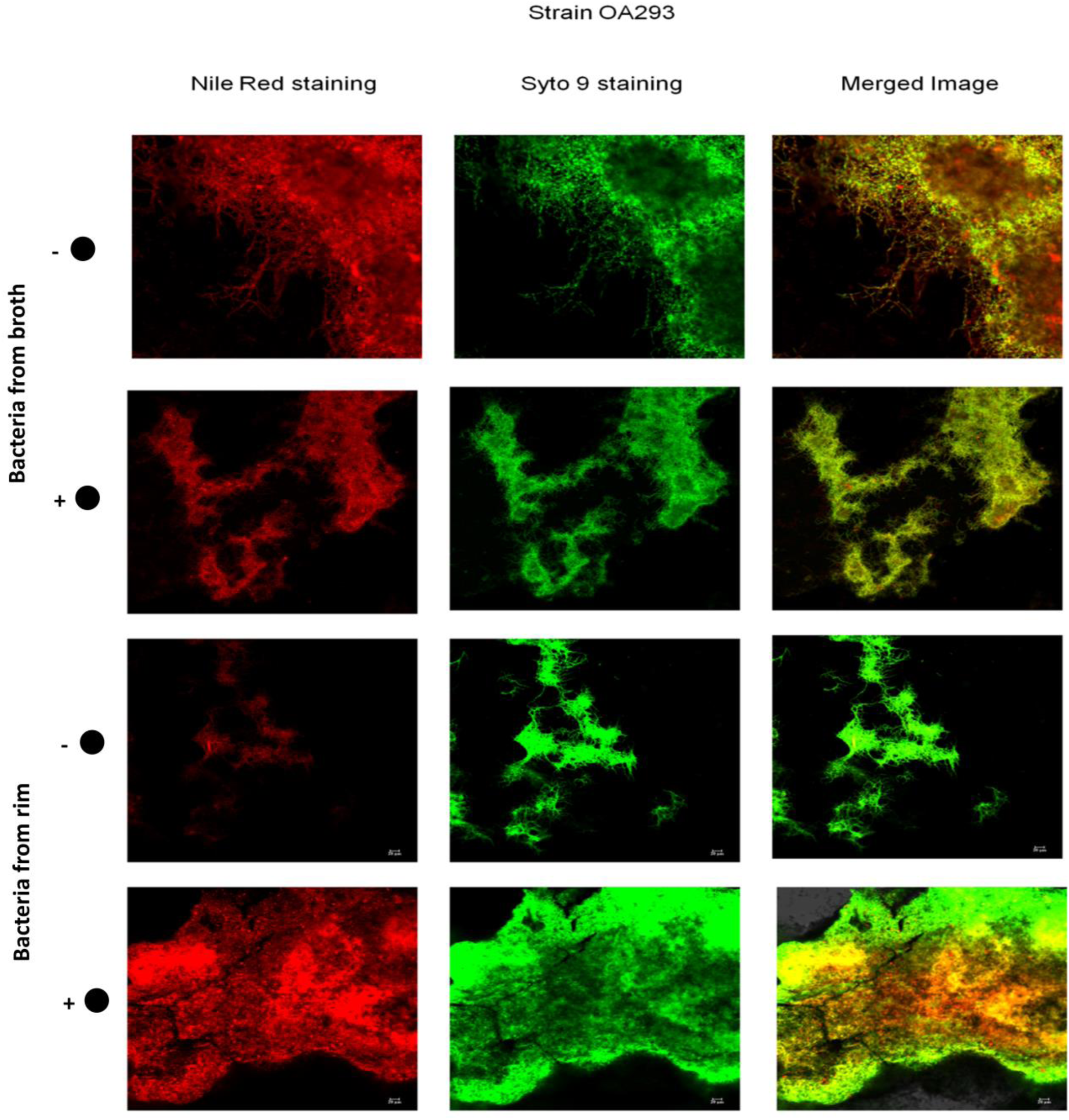
Confocal microscopy images show bacteria from broth and wall deposits. ‘+●’ and ‘-●’ represent bacteria from flasks with and without marbles, respectively. The differentiated aerial mycelium appears red and all live bacteria are shown green.

**Supplementary Figure 4:**
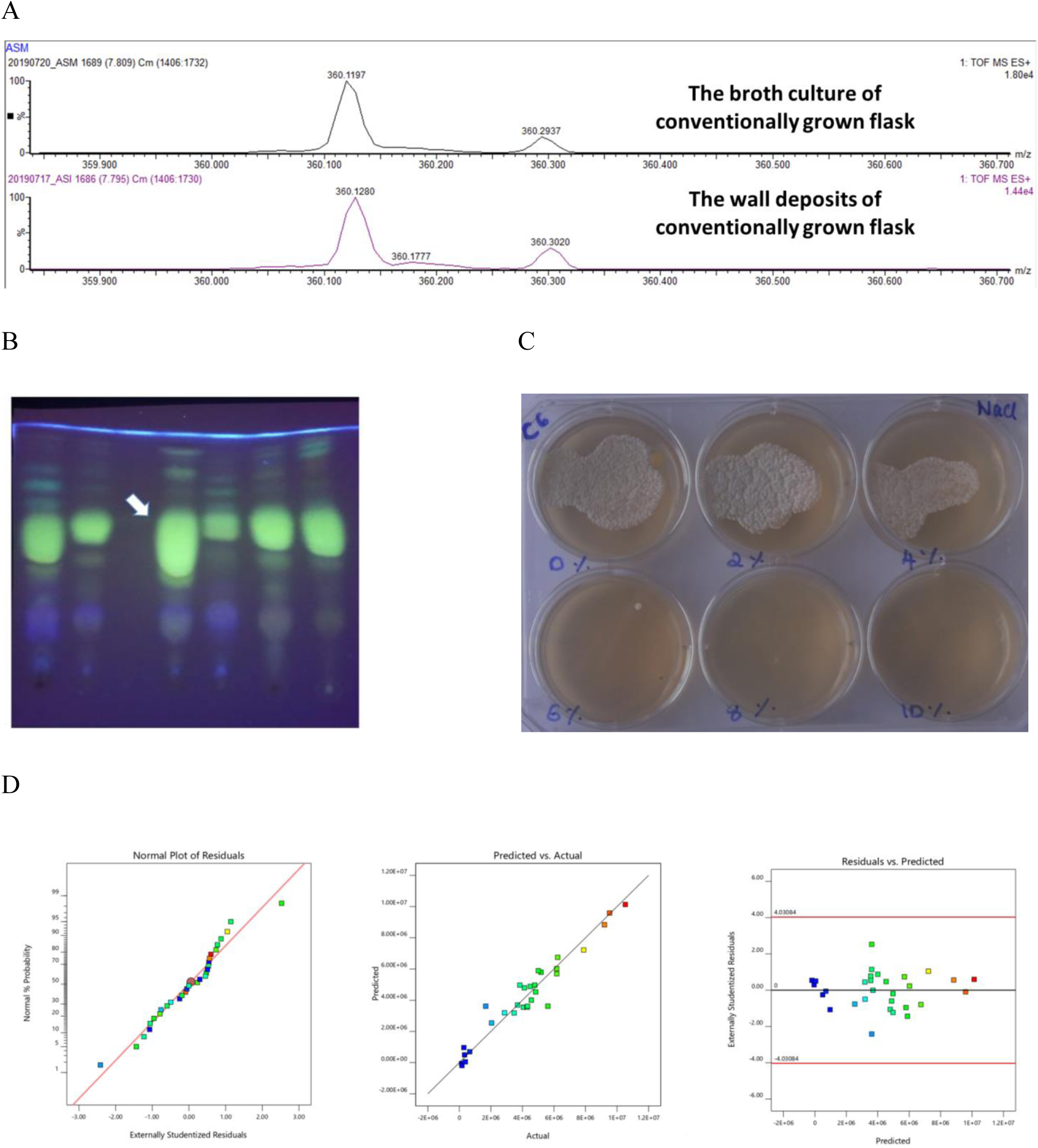
A. Measurement of trehalose in a culture without mechanical disruptors Trehalose was present in both wall deposits and the broth culture. B: Representative Thin layer chromatography plate from which chrysomycin was estimated. Arrow points at chrysomycins which are green fluorescent. C. Salt (NaCl) tolerance of *Streptomyces sp.*, OA161. It can tolerate upto 4 percent and thus a mild halophile. D. Diagnostic plots for Box-Behnken design

**Supplemetary Table 1.** shows the medium components and their respective concentrations used in the PBD design. A:D-Maltose; B:Yeast extract; C: Dextrose; D: Calcium carbonate; E:Temperature; F: Sodium chloride; G: Magnesium sulphate; H: Ferrous sulphate; J: Tween 80; K: Asparagine; L:Sodium dihydrogen phosphate; Y: Chrysomycins production.

|  | Factor<br>1 | Factor<br>2 | Factor<br>3 | Factor<br>4 | Factor<br>5 | Factor<br>6 | Factor<br>7 | Factor<br>8 | Factor<br>9 | Factor<br>10 | Factor<br>11 | Response |
| --- | --- | --- | --- | --- | --- | --- | --- | --- | --- | --- | --- | --- |
|  | A | B | C | D | E | F | G | H | J | K | L | Y |
| Run | g/L | g/L | g/L | g/L | C | % | g/L | mg/L | % | g/L | g/L | a.u |
| 1 | 5 | 0 | 2 | 2 | 28 | 2 | 2 | 0 | 0.1 | 5 | 2 | 1827693 |
| 2 | 10 | 2 | 6 | 1 | 32.5 | 1 | 1 | 5 | 0.05 | 2.5 | 1 | 2002728 |
| 3 | 10 | 2 | 6 | 1 | 32.5 | 1 | 1 | 5 | 0.05 | 2.5 | 1 | 2342932 |
| 5 | 5 | 0 | 10 | 0 | 37 | 2 | 0 | 10 | 0.1 | 5 | 0 | 575713.4 |
| 6 | 15 | 0 | 10 | 2 | 28 | 2 | 2 | 10 | 0 | 0 | 0 | 305544.2 |
| 7 | 5 | 4 | 10 | 0 | 37 | 2 | 2 | 0 | 0 | 0 | 2 | 2487039 |
| 8 | 10 | 2 | 6 | 1 | 32.5 | 1 | 1 | 5 | 0.05 | 2.5 | 1 | 3031723 |
| 9 | 15 | 4 | 10 | 0 | 28 | 0 | 2 | 0 | 0.1 | 5 | 0 | 7105.68 |
| 10 | 15 | 4 | 2 | 0 | 28 | 2 | 0 | 10 | 0.1 | 0 | 2 | 3435682 |
| 11 | 15 | 0 | 2 | 0 | 37 | 0 | 2 | 10 | 0 | 5 | 2 | 2380498 |
| 12 | 5 | 0 | 2 | 0 | 28 | 0 | 0 | 0 | 0 | 0 | 0 | 353246.4 |
| 13 | 5 | 4 | 10 | 2 | 28 | 0 | 0 | 10 | 0 | 5 | 2 | 7215017 |
| 14 | 10 | 2 | 6 | 1 | 32.5 | 1 | 1 | 5 | 0.05 | 2.5 | 1 | 2303496 |
| 15 | 15 | 4 | 2 | 2 | 37 | 2 | 0 | 0 | 0 | 5 | 0 | 3853949 |
| 16 | 5 | 4 | 2 | 2 | 37 | 0 | 2 | 10 | 0.1 | 0 | 0 | 3846336 |

**Supplementary Table 2.** shows Box-Behnken matrix of the experimental design along with the medium components with their concentrations.

|  |  | Factor 1 | Factor 2 | Factor 3 | Factor 4 | Response |
| --- | --- | --- | --- | --- | --- | --- |
| Block | Run | A:<br>D-Maltose<br>g/L | B:<br>Yeast extract<br>g/L | C:<br>Dextrose<br>g/L | D:<br>Calcium carbonate<br>g/L | Chrysomycins<br>a.u |
| Block 1 | 1 | 10 | 2 | 10 | 0 | 6.17649E+06 |
| Block 1 | 2 | 10 | 2 | 6 | 1 | 5.61584E+06 |
| Block 1 | 3 | 5 | 4 | 6 | 1 | 7.88417E+06 |
| Block 1 | 4 | 10 | 2 | 10 | 2 | 2.86747E+06 |
| Block 1 | 5 | 15 | 0 | 6 | 1 | 288238 |
| Block 1 | 6 | 10 | 2 | 6 | 1 | 1.67411E+06 |
| Block 1 | 7 | 5 | 0 | 6 | 1 | 330486 |
| Block 1 | 8 | 15 | 4 | 6 | 1 | 6.17558E+06 |
| Block 1 | 9 | 10 | 2 | 2 | 2 | 2.04669E+06 |
| Block 1 | 10 | 10 | 2 | 2 | 0 | 4.84351E+06 |
| Block 2 | 11 | 10 | 2 | 6 | 1 | 4.06793E+06 |
| Block 2 | 12 | 15 | 2 | 6 | 2 | 3.47532E+06 |
| Block 2 | 13 | 10 | 0 | 2 | 1 | 142894 |
| Block 2 | 14 | 10 | 4 | 2 | 1 | 4.12303E+06 |
| Block 2 | 15 | 10 | 2 | 6 | 1 | 4.29516E+06 |
| Block 2 | 16 | 10 | 0 | 10 | 1 | 167212 |
| Block 2 | 17 | 5 | 2 | 6 | 2 | 4.56646E+06 |
| Block 2 | 18 | 15 | 2 | 6 | 0 | 5.01052E+06 |
| Block 2 | 19 | 10 | 4 | 10 | 1 | 6.23688E+06 |
| Block 2 | 20 | 5 | 2 | 6 | 0 | 5.18363E+06 |
| Block 3 | 21 | 15 | 2 | 2 | 1 | 4.32849E+06 |
| Block 3 | 22 | 5 | 2 | 10 | 1 | 4.49389E+06 |
| Block 3 | 23 | 10 | 2 | 6 | 1 | 3.83515E+06 |
| Block 3 | 24 | 10 | 4 | 6 | 2 | 3.69376E+06 |
| Block 3 | 25 | 10 | 4 | 6 | 0 | 9.20238E+06 |
| Block 3 | 26 | 10 | 0 | 6 | 2 | 659148 |
| Block 3 | 27 | 15 | 2 | 10 | 1 | 1.05265E+07 |
| Block 3 | 28 | 10 | 0 | 6 | 0 | 382622 |
| Block 3 | 29 | 5 | 2 | 2 | 1 | 9.5291E+06 |
| Block 3 | 30 | 10 | 2 | 6 | 1 | 4.78289E+06 |

## Notes

### Competing Interest Statement

The authors have declared no competing interest.

